# First report of emerging snake fungal disease caused by *Ophidiomyces ophiodiicola* from Asia in imported captive snakes in Japan

**DOI:** 10.1101/2020.09.03.281154

**Authors:** Yoshinori Takami, Yumi Une, Ikki Mitsui, Chizuka Hemmi, Youki Takaki, Tsuyoshi Hosoya, Kyung-Ok Nam

## Abstract

Snake fungal disease (SFD) (Ophidiomycosis) is an emerging infectious disease caused by the fungus *Ophidiomyces ophiodiicola* which has been affecting wild and captive snakes in North America, Europe, and Australia. We report the cases of 12 imported captive colubrid snakes in Japan suspected of having SFD. Pathological and microbiological examinations were performed, and the results confirmed the diagnosis of SFD in two snakes, which indicated that the remaining sympatrically raised snakes also had SFD since they exhibited similar lesions. The oral administration of ciprofloxacin in addition to itraconazole had a significant treatment effect. This is the first report of SFD in Asia caused by *O. ophiodiicola*. To prevent the expansion of SFD in the natural environment in Japan, there is a need to evaluate the SFD carrier status of imported snakes, the pathogenicity of the infection in native snakes, and the prevalence and distribution of SFD in wild and captive snakes. Measures also must be taken to prevent endemicity globally.

## Introduction

Chytridiomycosis in amphibians, white-nose syndrome in bats, and snake fungal disease (SFD) (Ophidiomycosis) have been known as serious fungal diseases that harm ecosystems^1–4^. SFD is caused by the fungus *Ophidiomyces ophiodiicola* (*O. ophiodiicola*) 5]. Since 2006, severe skin infections have been reported in association with a decline in timber rattlesnake (*Crotalus horridus*) populations in the northeastern United States^1^. In 2008, a similar infection involving a fungal pathogen occurred in the endangered species massasaugas (*Sistrurus catenatus*) in Illinois, USA^5^. The infection became known as SFD, and by 2015 SFD was reported in almost every wild snake in the eastern United States^6^.

Examinations of isolates reveal that *O. ophiodiicola* has been present in captive snakes in the eastern United States since 1986^7^. In contrast, isolates of wild snakes were not reported until 2008^8^. Therefore, *O. ophiodiicola* is thought to have been spread from the captive to the wild snake population^5^.

Currently the known geographical distribution of *O. ophiodiicola* is wider for captive snakes than wild snakes^5^. In the United States, isolates have been obtained from captive snakes in California, Georgia, Maryland, New Mexico, New York, and Wisconsin^7^. Outside North America, *O. ophiodiicola* has been isolated from lesions of captive snakes in the United Kingdom, Germany and Australia^7,9^. To date, SFD has been documented in more than 30 snake species^10^ and is an emerging infectious disease affecting wild and captive snakes throughout North America, Europe, and Australia^10^; however so far there are no reports from Asia.

In this study, we report the first epidemic case in Asia caused by *O. ophiodiicola* in imported captive snakes.

## Results

The owner of 12 colubrid snakes (one hognose snake [*Heterodon sp*.], one California kingsnake [*Lampropeltis getula californiae*], two Mexican black kingsnakes [*Lampropeltis getula nigrita*], one green vine snake [*Oxybelis fulgidus*], three corn snakes [*Pantherophis guttatus*], two black rat snakes [*Pantherophis obsoletus*], and two Texas rat snakes [*Pantherophis obsoletus lindheimeri*]) noticed blisters, pustules, and crusts on the snakes’ skins, and had taken them to a veterinary hospital from April to October 2019 (snakes are numbered 1 to 12 in Table 1). The first snake which presented SFD was a wild-caught green vine snake (No. 1) that was purchased on April 21, 2019. Subsequently, the remaining 11 captive snakes successively presented skin lesions resembling SFD between June and October of 2019. Eventually, five of the 12 snakes (No. 1-5) died; among them, three (No. 2, 4, 5) were pathologically and microbiologically examined. The five snakes that died received only antibiotics (ciprofloxacin; 5 mg/kg, orally, once a day). In contrast, the seven surviving snakes (No. 6-12) received antifungal itraconazole (5 mg/kg, orally, once a day) in addition to the same dosage of ciprofloxacin. The skin lesions on these seven snakes disappeared macroscopically within 14–37 days after initiation of treatment (mean: 32.4 days); although the skin lesions recurred in one snake (No. 6) on day 57 (Table 1 summarizes these results).

**Table 1.**
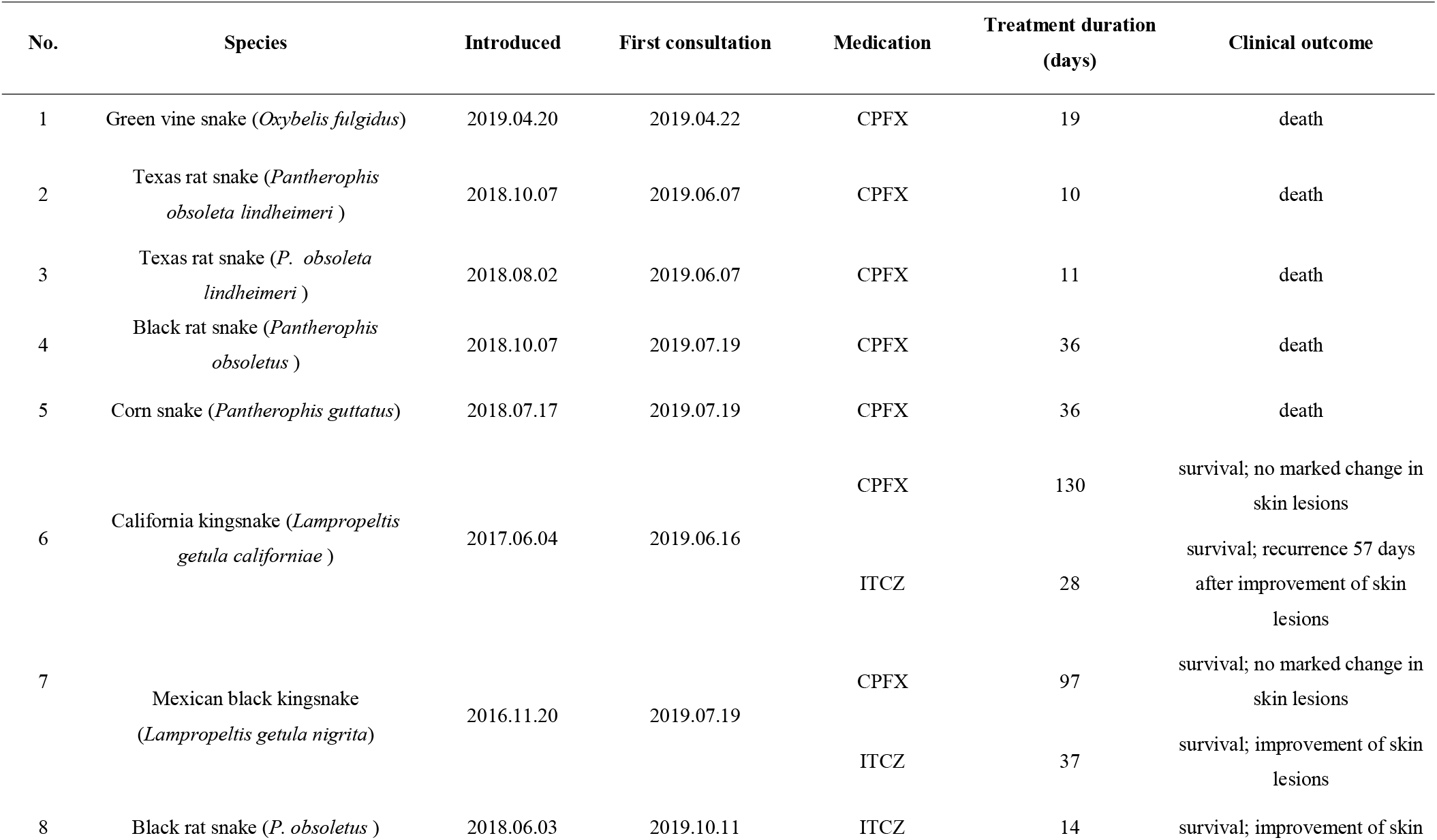
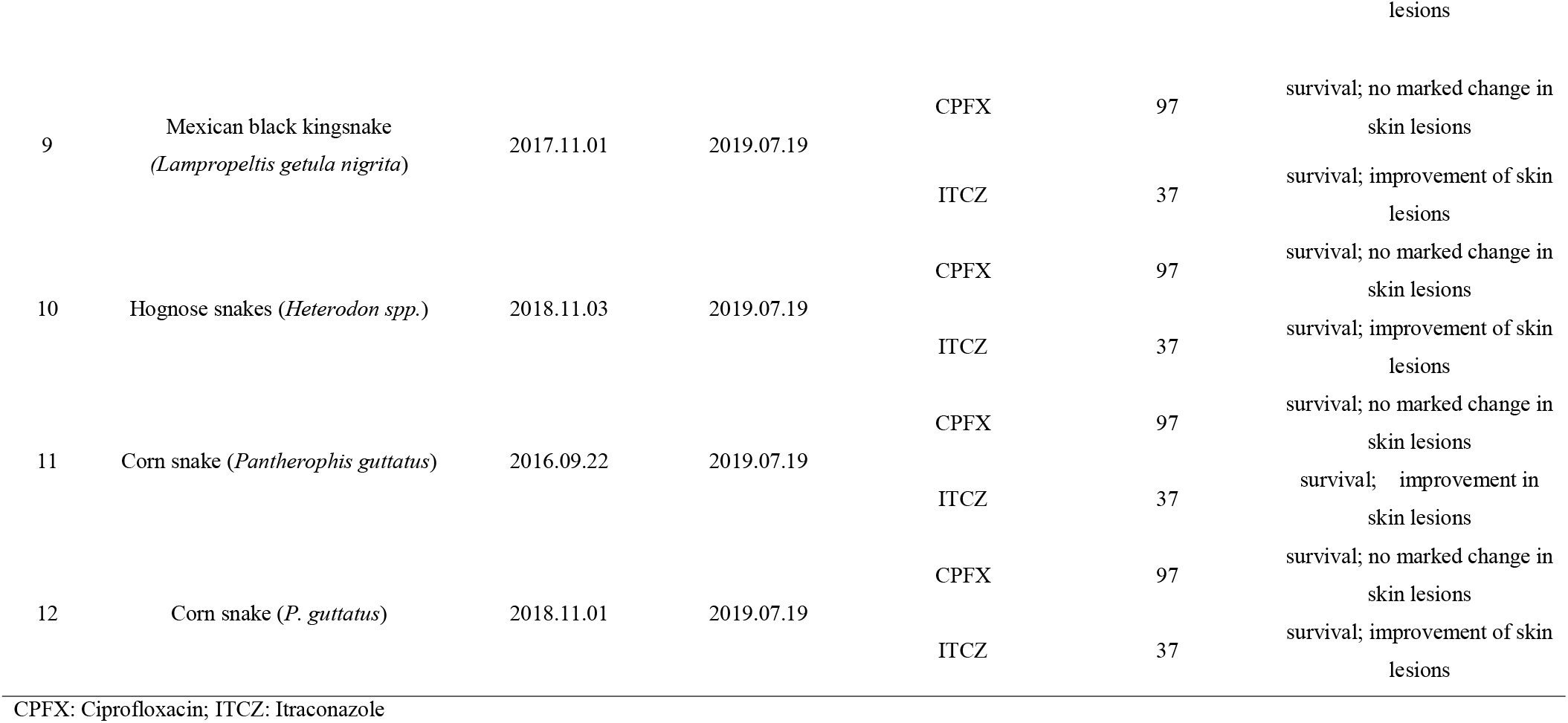
Colubrid snake species, date of introduction, date of first consultation, medication, treatment duration, and clinical outcome.

Skin changes were uniform in all snakes; specifically, they had discolouration (yellow, light brown, or red), turbidity, increased fragility, and scale thickening, which occurred in widespread bands and patches throughout their trunks.

The skins were easily exfoliated, and multifocal crust and blister formation were observed throughout their bodies. Pathological examination of the skin lesions on snakes No. 2, 4, and 5 revealed severe degeneration and scale tissue necrosis accompanied by a high degree of crusting where the fungus were observed (Figure 1 A-G). No significant changes in the internal organs were recognized.

**Figure 1.**
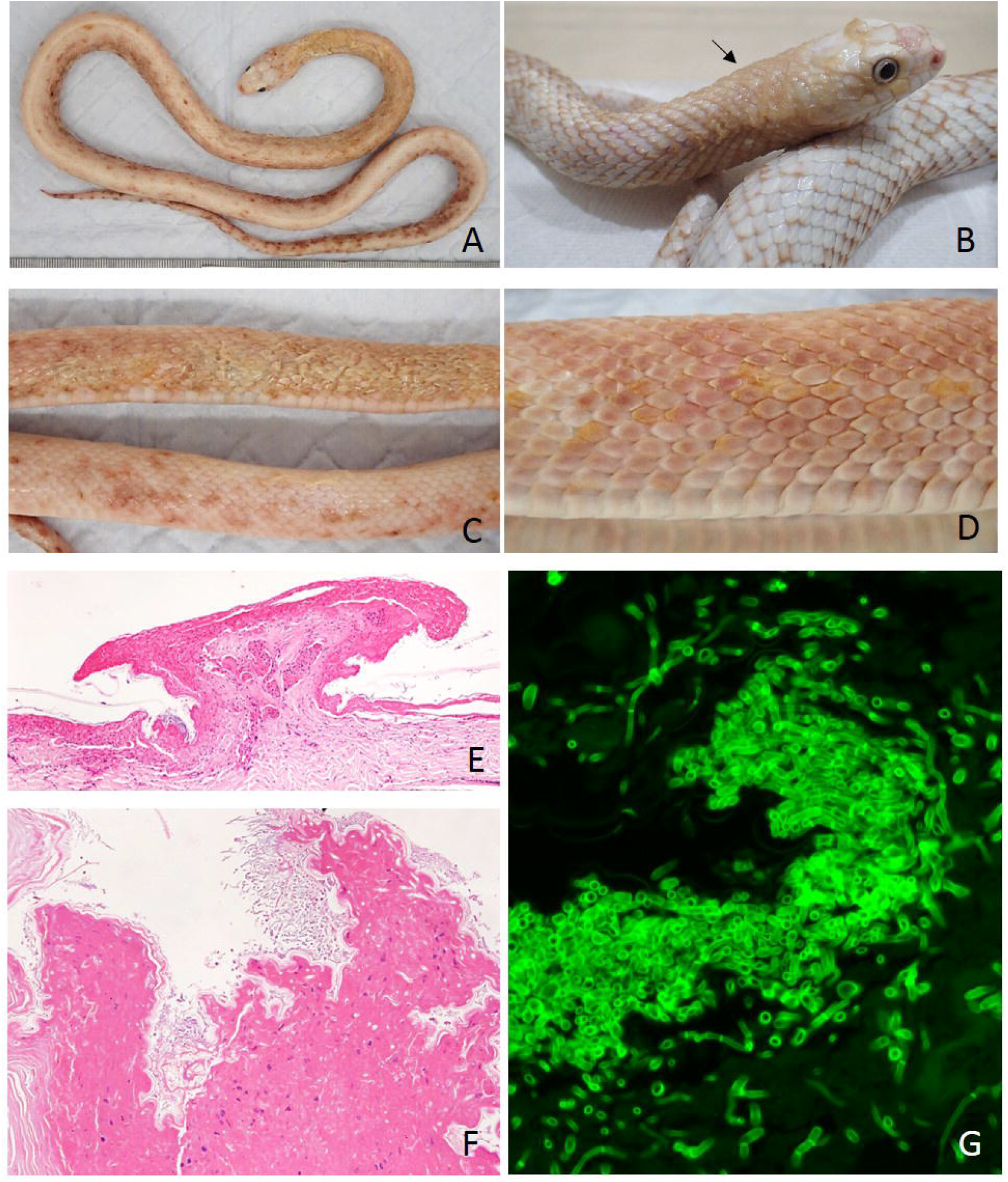
Pathological features of ophidiomycosis. **A**. No. 4 (*Pantherophis obsoletus*). The skin of the cranial body is extensively necrotic, and the skin of the caudal body has multifocal bleeding and crusting lesions. They are severe at the tail. **B**. Exterior view prior to death of No. 2 (*P. obsoletus lindheimeri*). Discolouration of scale (browning); at first, the edges of the individual scales change colour, then gradually the whole scale is involved. As the discolouration progresses further, it becomes more diffuse as in the skin on the neck (arrow). **C**. The skin lesions of No. 4 (*P. obsoletus*). Upper: extensive necrosis of skin; Lower: multifocal hyperaemia and crusting. **D**. No. 5 (*P. guttatus*). Necrotic scales are yellow. **E**. The skin lesions (Haematoxylin and eosin stain) of No. 5 (*P. guttatus*). The entire scale is necrotic; fungal growth foci are seen at the base of the scale (arrows). F: The skin lesions (Haematoxylin and eosin stain) of No.5 (*P. guttatus*). High magnification of fungal growth foci with severe necrosis of the scales (arrows). **G**. Fungi in tissue (Fungiflora Y stain) of No. 4 (*P. obsoletus*). Crust and fungal growth are present; fungiflora Y staining shows that the fungal cell membrane is fluorescent green. The fungus is penetrating deeply into the surface and crust. The fungus is rather thin and has a septum

In the three subjects (No, 2, 4, and 5) we examined, the fungus occurred on No. 2 and 5. We obtained several isolates by isolating spores from the powdery colonies formed on the snake tissues and inoculating them on an agar plate. After isolation, we confirmed that all the colonies had identical morphology and chose one representative from each sample (NBRC 114438 from *P. obsoletus lindheimeri* [No. 2] and NBRC 114439 from *P. obsoletus* [No. 5]). We confirmed that the micromorphology for the conidia-producing structure agreed with the previous descriptions^7,11^ (Figure 2).

**Figure 2.**
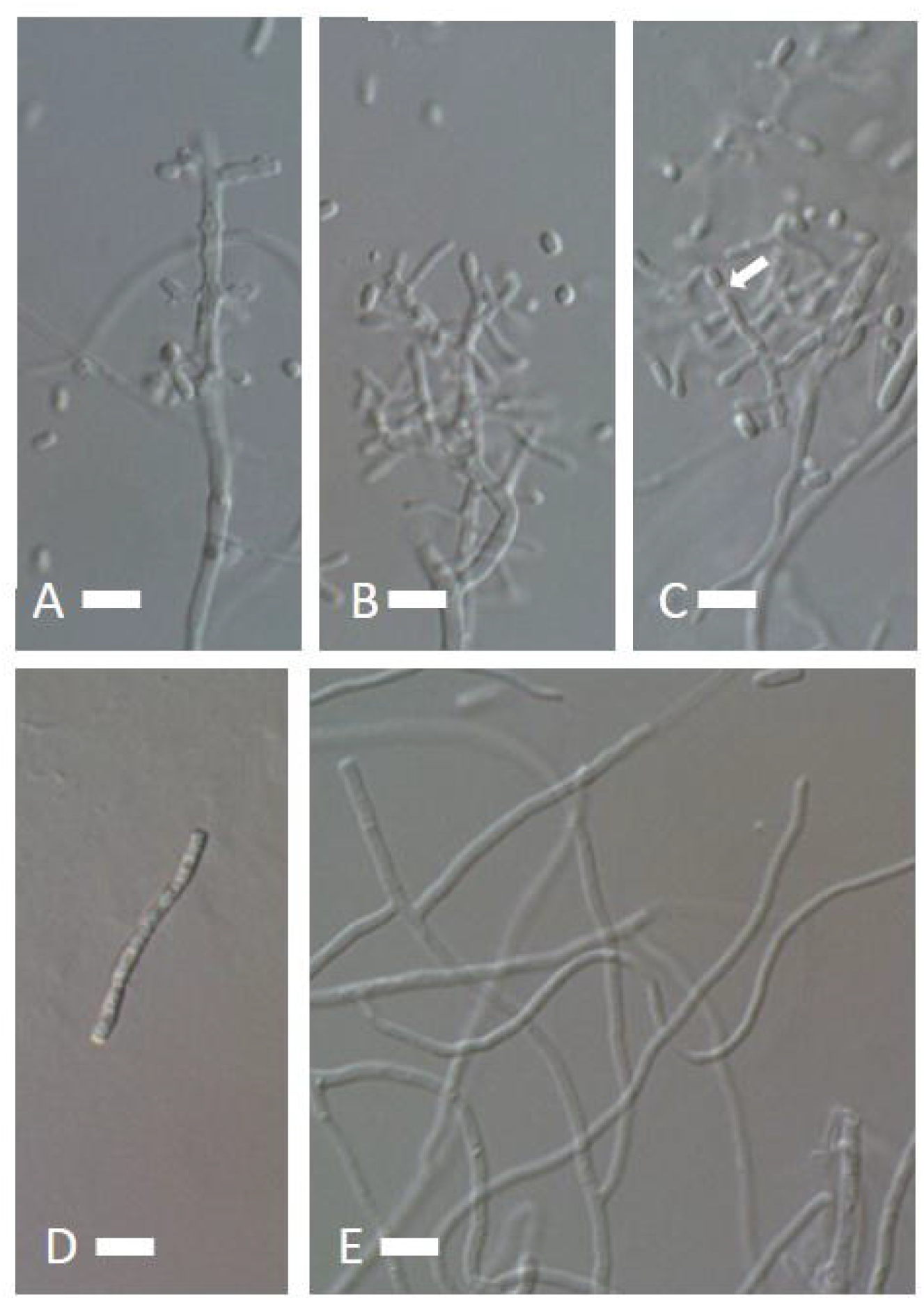
Micrograph of *Ophidiomyces ophiodiicola*. **A:** Vestiges of conidia attachment in conidia-producing hyphae. **B:** Conidia produced laterally from the conidia-producing hyphae. **C:** Thallic conidia with separating cell (arrow). **D:** Arthroconidia produced in a chain, detached from the hyphae. **E:** Undulating hyphae. Scale: 10 μm.

The obtained sequences (LC521891 from NBRC 114438 and LC521892 from NBRC 114439) were identical, and were searched using Basic Local Alignment Search Tool (BLAST) and compared to the sequence database in The National Centre for Biotechnology Information (NCBI). The most similar sequences with E value = 0.0 were all for *Ophidiomyces ophiodiicola* except for EU715819 (*Chrysosporium* sp. = *O. ophiodiicola*). The sequence with the highest similarity was KF225599 (*O. ophiodiicola* voucher MYCO-ARIZ AN0400001) with 99.52% identity with 100% query coverage, with the rest showing a 98.13–100 % identity. These analyses confirmed that the isolated fungus was *O. ophiodiicola*, also known as O. ophiicola (http://www.indexfungorum.org/names/NamesRecord.asp?RecordID=545168).

## Discussion

Our examination confirmed an SFD diagnosis in two snakes, and the remaining sympatrically raised snakes had exhibited similar lesions. *Ophidiomyces ophiodiicola* was initially classified as *Chrysosporium* which is widely known as an asexual state of the order Onygenales, and later transferred to the monotypic genus *Ophidiomyces* based on the molecular phylogeny^7^. It has a high pathogenicity, with a mortality rate of ≥ 90% in some snake species such as massasaugas in the eastern United States^12^. There are many questions raised about this disease, in particular, regarding the effects of the disease on wild snakes and captive snakes, the presence of infection vectors, and the change in prevalence over time^13^.

According to the owner, the present outbreak among the snakes began after the introduction of a wild-caught green vine snake (*Oxybelis fulgidus*) indigenous to South America. Because the obtained sequences (LC521891 from NBRC 114438 and LC521892 from NBRC 114439) were identical, it is considered that a single isolate caused the present outbreak case. Based on these results, *O. fulgidus* was the suspected infection source. Since no case of SFD has been reported previously in *O. fulgidus*, this snake seems to be a new host and the fungus may have a wider host range than previously suspected.

There are 36 wild snake species in Japan with high diversity in the mid-latitude areas. Those snakes occupy an important position as predators in the food chain. To prevent expansion of SFD into the natural environment in Japan, there is a need to evaluate the SFD carrier status of imported snakes, the pathogenicity of the infection in native snakes, and the clarification of prevalence and distribution of SFD in wild and captive snakes.

## Methods

### Isolation of the pathogen

For fungal isolation, three pieces of 10 x 5 x 5 mm tissue samples were collected from snakes No. 2, 4, and 5 were inoculated on a corn meal agar plate (CMA, Nissui, Tokyo), and incubated at room temperature for approximately two weeks to allow colonization of the fungus. White, compact, powdery fungal colonies appeared on two samples, which were associated with bacteria and faster-growing fungi. To isolate the target fungus, we used Skerman’s micromanipulator (Skerman, VBD 1968)^14^, which allows for single spore isolation. For microscopic observation of the morphology, potato dextrose agar plates (PDA, Nissui, Tokyo) were inoculated with the isolate in the centre and incubated at 25 C° for 10 days. The isolates were deposited to the Biological Resource Centre, National Institute of Technology and Evaluation (NITE BRC, Tokyo, Japan). The tissue samples were collected from dead snakes brought to a veterinary hospital and therefore ethical approval was not required from the Institutional Review Board.

### Culture and sequencing

The isolates were cultured in 2 ml of 2% malt extract (Difco). DNA extraction and polymerase chain reaction (PCR) using primers ITS1F and ITS4^15^ for the ITS-5.8S sequence region were performed as described by Itagaki et al.^16^.

### Pathological examination

All tissues were fixed in 10% neutral buffered formalin. Paraffin sections (3-4 μm thick) were stained with haematoxylin and eosin, periodic acid Schiff, and Fungiflora Y (Cosmo Bio CO., LTD. Japan).

## Data Availability

All data generated or analysed during this study are included in this published article.

## Acknowledgements

This work was supported by Japan Society for the Promotion of Science KAKENHI Grant-in-Aid for Scientific Research (C) (grant number 18K05692).

## Authors’ Contributions

YU, Y Takami, and TH performed post-mortem examinations and coordinated diagnostic testing; Y Takaki assisted with sample collection; YU, IM, and CH performed histological examinations; TH and KON conducted sequence characterisation and fungal culture characterisation; YU Y Takami, and TH led the manuscript preparation. Y Takami and YU contributed equally to this manuscript. All authors contributed to writing the manuscript and approved the final version.

